# Comparative genomics confirms a rare melioidosis human-to-human transmission event and reveals incorrect phylogenomic reconstruction due to polyclonality

**DOI:** 10.1101/804344

**Authors:** Ammar Aziz, Bart J. Currie, Mark Mayo, Derek S. Sarovich, Erin P. Price

**Affiliations:** Global and Tropical Health Division, Menzies School of Health Research, Charles Darwin University, Darwin, NT, Australia; Infectious Diseases Department, Royal Darwin Hospital, Darwin, NT, Australia; GeneCology Research Centre, University of the Sunshine Coast, Sippy Downs, QLD, Australia; Sunshine Coast Health Institute, Birtinya, QLD, Australia

**Keywords:** human-to-human transmission, *Burkholderia pseudomallei*, Melioidosis, phylogenomics, comparative genomics, strain mixtures, bioinformatics

## Abstract

Human-to-human transmission of the melioidosis bacterium, *Burkholderia pseudomallei*, is exceedingly rare, with only a handful of suspected cases documented to date. Here, we used whole-genome sequencing (WGS) to characterise one such unusual *B. pseudomallei* transmission event, which occurred between a breastfeeding mother with mastitis and her child. Two strains corresponding to multilocus sequence types (STs) 259 and 261 were identified in the mother’s sputum from both the primary culture sweep and in purified colonies, confirming an unusual polyclonal infection in this patient. In contrast, primary culture sweeps of the mother’s breast milk and the child’s cerebrospinal fluid and blood samples contained only ST-259, indicating monoclonal transmission to the child. Analysis of purified ST-259 isolates showed no genetic variation between mother and baby isolates, providing the strongest possible evidence of *B. pseudomallei* transmission, probably via breastfeeding. Next, phylogenomic analysis of all isolates, including the mother’s mixed ST-259/261 sputum sample was performed to investigate the effects of mixtures on phylogenetic inference. Inclusion of this mixture caused a dramatic reduction in the number of informative SNPs, resulting in branch collapse of ST-259 and ST-261 isolates, and several instances of incorrect topology in a global *B. pseudomallei* phylogeny, resulting in phylogenetic incongruence. Although phylogenomics can provide clues about the presence of mixtures within WGS datasets, our results demonstrate that this methodology can lead to phylogenetic misinterpretation if mixed genomes are not correctly identified and omitted. Using current bioinformatic tools, we demonstrate a robust method for bacterial mixture identification and strain parsing that avoids these pitfalls.

**Impact Statement:** *Burkholderia pseudomallei* is the causative agent of melioidosis, a tropical disease of high mortality. *B. pseudomallei* infections occur almost exclusively through contact with contaminated soil and water. Using whole-genome sequencing (WGS), we investigated a rare case of suspected *B. pseudomallei* transmission from mother to child. The mother’s sputum, breast milk and the baby’s blood and cerebrospinal fluid (CSF) specimens were collected, and DNA was extracted from both pure colonies and primary culture sweeps to capture potential strain mixtures. In-depth analysis of genetic variants identified two strains in the mother’s sputum belonging to multilocus sequence types ST-259 and ST-261, whereas the child was infected with only ST-259. Comparative genomics revealed no genetic differences between mother and child ST-259 isolates, providing the strongest possible evidence of transmission to the child via breast milk. The sputum strain mixture was subsequently used to develop a bioinformatic method for identification and quantification of mixtures from WGS data. Using this method, we found ST-259 and ST-261 at an 87%:13% ratio, respectively. Finally, we demonstrate the negative impact that even a single strain mixture event can have on both within-ST and global phylogenomic inferences. Our findings highlight the need for bioinformatic quality control to avoid unintended consequences of phylogenomic incongruence and branch collapse.

**Data Summary:**

1. Whole-genome sequencing data have been deposited in the NCBI Sequence Read Archive (SRA) and GenBank under BioProject accession number <u>PRJNA559002</u>.
2. The GenBank accession number for MSHR0643 assembly is VXLH00000000.1.
3. The SRA accession numbers for all raw sequence data are listed in Table 1.

**Repositories:** All sequencing data generated as part of this study can be found under the NCBI BioProject PRJNA559002 with accession numbers listed in Table 1.

## Introduction

*Burkholderia pseudomallei*, a Gram-negative environmental bacterium found in soil and water in mostly tropical regions, is the causative agent of melioidosis [1]. This underreported and historically neglected disease is being increasingly recognised to be endemic in diverse tropical regions globally, and hyperendemic in northern Australia and Southeast Asia [2]. *B. pseudomallei* is an opportunistic bacterium that most commonly affects people who are in regular contact with soil and water, with percutaneous inoculation and inhalation the main routes of infection, with infection by ingestion uncommon [1, 3]. The high mortality rate of melioidosis (10-40%) even with antibiotic treatment [4], combined with the intrinsic resistance of *B. pseudomallei* against a wide range of antibiotics [5], highlight the significant public health importance of this bacterium [1]. Increasing awareness and detection of melioidosis in new locales and the lack of a vaccine towards *B. pseudomallei* have further increased the global public health significance of this pathogen [6]. Due to these factors, *B. pseudomallei* is considered a Tier 1 Select Agent pathogen due to its potential for misuse as a biological warfare agent [7].

Multilocus sequence typing (MLST) is a commonly used genotyping method for determining the population structure, geography, source attribution and transmission patterns of many bacterial pathogens, including *B. pseudomallei* [8]. With the advent of whole-genome sequencing (WGS), simultaneous genomic characterisation, phylogeography, multilocus sequence type (ST) determination, antibiotic resistance profiling and fine-scale resolution of *B. pseudomallei* population structure, evolution and transmission profiles have become possible [9]. WGS has also assisted with the identification of polyclonal *B. pseudomallei* infections, including one reported instance of a polyclonal infection with the same ST [10].

Although rare, a handful of suspected cases of human-to-human *B. pseudomallei* transmission have been documented, including between siblings with cystic fibrosis [11], between siblings with diabetes [12], between an American Vietnam veteran diagnosed with *B. pseudomallei*-associated prostatitis and his spouse, but supported only by serology [13], and three cases between mother and child [3, 14]. In one of the mother-to-child transmission cases, a mother with *B. pseudomallei*-associated mastitis in her left breast was suspected to have transmitted this pathogen to her breastfed infant [3]. Mother-to-child *B. pseudomallei* transmission via transplacental, breast, or perinatal routes has been suspected in a handful of other human cases [3, 14, 15], and in animals [16]. However, no human-to-human transmissions reported to date have been confirmed using WGS, which is essential for ruling out concomitant environmental sources of infection. In the current study, WGS was used to understand the dynamics of this unusual human-to-human transmission event, which was also characterised by a polyclonal infection detected in the mother’s sputum. Using comparative genomics, we provide the strongest possible evidence for human-to-human *B. pseudomallei* transmission between mother and child. We next examined the impact of the strain mixture identified in the mother’s sputum sample on phylogenetic interpretations of maternal to child transmission. We observed confounding phylogenomic results when the single mixed genome was included in the analysis, a finding that has implications for fine-scale phylogenomic investigation of outbreak, source attribution, or host transmission studies.

## Methods

### Case history and bacterial culture

The clinical history of the mother-to-child transmission case has been described elsewhere [3]. Briefly, a seven-month old breast-feeding child from a remote region in northern Australia was hospitalised in 2003 with acute cough, fever and tachypnoea. During this admission, the mother was observed to have a fever and pleuritic chest pain and was subsequently diagnosed with mastitis in the left breast. Upon *B. pseudomallei* culture confirmation in the child’s cerebrospinal fluid (CSF), blood, and nasal and throat swabs, the mother was also tested for *B. pseudomallei* infection in blood, sputum, and multiple breast milk specimens, and from nasal, throat, and rectal swabs. Of these, *B. pseudomallei* was isolated from the mother’s breast milk and sputum (Table 1). All clinical specimens were cultured onto Ashdown’s media as described elsewhere [17]. DNA extractions were performed [18] on a sweep of the primary culture streak (herein referred to as primary culture sweeps) of each *B. pseudomallei*-positive clinical specimen in an effort to capture potential strain mixtures in these original specimens, and subsequently from individually purified colonies derived from these specimens.

**Table 1.** Summary of ST-259 and ST-261 *Burkholderia pseudomallei* isolates.

| Isolate ID* | Sample Type | Patient | Multilocus sequence type | NCBI Accession Numbers |
| --- | --- | --- | --- | --- |
| MSHR1574 | CSF | Child | ST-259 | SRR9959037 |
| MSHR1574_Sweep | CSF | Child | ST-259 | SRR9959038 |
| MSHR1580 | Blood | Child | ST-259 | SRR9959039 |
| MSHR1580_Sweep | Blood | Child | ST-259 | SRR9959040 |
| MSHR1583 | Breast milk | Mother | ST-259 | SRR9959042 |
| MSHR1583_Sweep | Breast milk | Mother | ST-259 | SRR9959036 |
| MSHR1631 | Sputum | Mother | ST-259 | SRR9959045 |
| MSHR1631_Mixed | Sputum | Mother | ST-259 and ST-261 | SRR9959043 |
| MSHR1581 | Sputum | Mother | ST-261 | SRR9959044 |
| MSHR1581_Sweep | Sputum | Mother | ST-261 | SRR9959041 |
| MSHR0120 | Blood | Other<sup>#</sup> | ST-259 | SRX1465234 |
| MSHR0669 | Blood | Other<sup>#</sup> | ST-259 | SRR9959034 |
| MSHR1224 | Blood | Other<sup>#</sup> | ST-259 | SRR9959035 |
| MSHR1328 | Sputum | Other<sup>#</sup> | ST-259 | SRR10134765 |
| MSHR1357 | Abscess | Other<sup>#</sup> | ST-259 | SRR10134764 |
| MSHR3509 | Blood | Other<sup>#</sup> | ST-259 | SRR10134763 |
| MSHR0643 | Urine | Other<sup>#</sup> | ST-259 | SRR9959033 |

\*Isolates with the “\_Sweep” suffix were obtained from primary culture sweeps to capture *B. pseudomallei* population diversity. Of these, MSHR1631\_Mixed was the only sample found to contain a mixture of two genotypes. Isolates without the “\_Sweep” suffix were obtained from purified single colonies derived from the “\_Sweep” culture. <sup>#</sup>Temporally and geographically distinct clinical ST-259 isolates obtained between 1992 and 2009 from other patients living in the Top End region of the Northern Territory.

### Whole-genome sequencing and *in silico* MLST

As part of the ongoing Darwin Prospective Melioidosis Study (DPMS), which commenced in 1989 [19], all mother and child primary culture sweeps and purified colonies (i.e. isolates) were subjected to WGS using the Illumina HiSeq2500 platform (Australian Genome Research Facility, Melbourne, Australia). WGS was performed on primary culture sweeps and isolates from the mother’s (*n*=6) and child’s (*n*=4) specimens. Reference-assisted draft genome assemblies were performed using MGAP v1.0 (default settings) [20], with the closed Australian MSHR1153 genome (CP009271.1 and CP009272.1 for chromosomes 1 and 2, respectively) [21] as reference. *In silico* MLST was performed using the PubMLST *B. pseudomallei* database available at http://pubmlst.org/bpseudomallei/ [22]. For the mixed-strain sample (MSHR1631_Mixed) manual allele assignment was performed by inspecting alignment files using Tablet [23] and parsing single-nucleotide polymorphisms (SNPs) corresponding to the two strains based on allele abundance.

### Comparative genomics and mixture analysis

Comparative genomic analysis was performed with the default settings of SPANDx v3.2 (https://github.com/dsarov/SPANDx) [24], which wraps Burrows-Wheeler Aligner [25], SAMtools [26], the Genome Analysis Toolkit (GATK v3.2.2) [27], BEDTools [28], and SNPEff [29] into a single pipeline. Mapping was carried out using the closed Australian genome MSHR1153 [21] as the reference, with the SPANDx *-i* flag enabled to provide insertion-deletion (indel) variant identification. Heterozygous SNPs in each isolate were enumerated from GATK UnifiedGenotyper VCF output. One sweep culture, MSHR1631_Mixed, exhibited a substantial number of heterozygous SNPs when compared with all other isolates and sweep cultures, so was further investigated as a possible mixture. Variant identification in MSHR1631_Mixed was determined using GATK v4.1 HaplotypeCaller [30] due to its ability to natively handle polyploid samples. Variant filtering was performed using the parameters described in SPANDx v3.2 [24]. For each heterozygous SNP identified in MSHR1631_Mixed, the depth (number of reads) supporting each allele was extracted from the VCF file and normalised by the total read depth at that SNP position. Additionally, to ensure robust variant calling and to assess mixture composition, we tested multiple ploidy settings (*n* = 2, 3, 4, and 5).

### Phylogenomic analysis

A maximum parsimony (MP) phylogenetic tree representing a global snapshot of *B. pseudomallei* isolates was constructed using orthologous, biallelic, core-genome SNPs identified across 145 publicly available genomes [31], which included the ten new isolates/sweep cultures sequenced as part of this study. To investigate *B. pseudomallei* transmission from mother to child, a combined SNP-indel [32] MP tree was constructed using all ST-259 isolates, with the ST-259 genome MSHR0643 as reference. MSHR0643 was chosen as the reference genome as it had the fewest contigs of any ST-259 strain (*n*=93). Also included were seven additional temporally distinct ST-259 isolates (Table 1). MP phylogenetic tree construction and bootstrapping (300 replicates) were performed using PAUP* v4.0a165 and visualised with iTOL v4 [33].

### Pulsed-field gel electrophoresis (PFGE)

*Spe*I DNA-digested PFGE was performed on mother and child isolates as previously described [34].

## Results and Discussion

*B. pseudomallei* causes melioidosis, a life-threatening disease with a predicted global incidence of ∼165,000 cases annually [2]. Almost all *B. pseudomallei* infections occur via contact with contaminated water or soil, while human-to-human transmission events are exceedingly rare [35]. Here, we used genomics to examine, in high resolution, one such human-to-human transmission event where a nursing mother with culture-confirmed melioidosis mastitis was suspected to have transmitted *B. pseudomallei* to her child through contaminated breast milk [3]. PFGE analysis on isolates retrieved from the mother and her child shortly after diagnosis identified two pulsotypes in the mother’s sputum isolates (Figure 1), suggesting a potential polyclonal infection. To further understand this unusual case, WGS was performed on all available specimens from these cases to elucidate transmission dynamics from mother to child, to investigate the potential presence of within-host strain mixtures in the mother, and finally, to examine the effects of strain mixtures on downstream phylogenomic interpretations.

**Figure 1.**
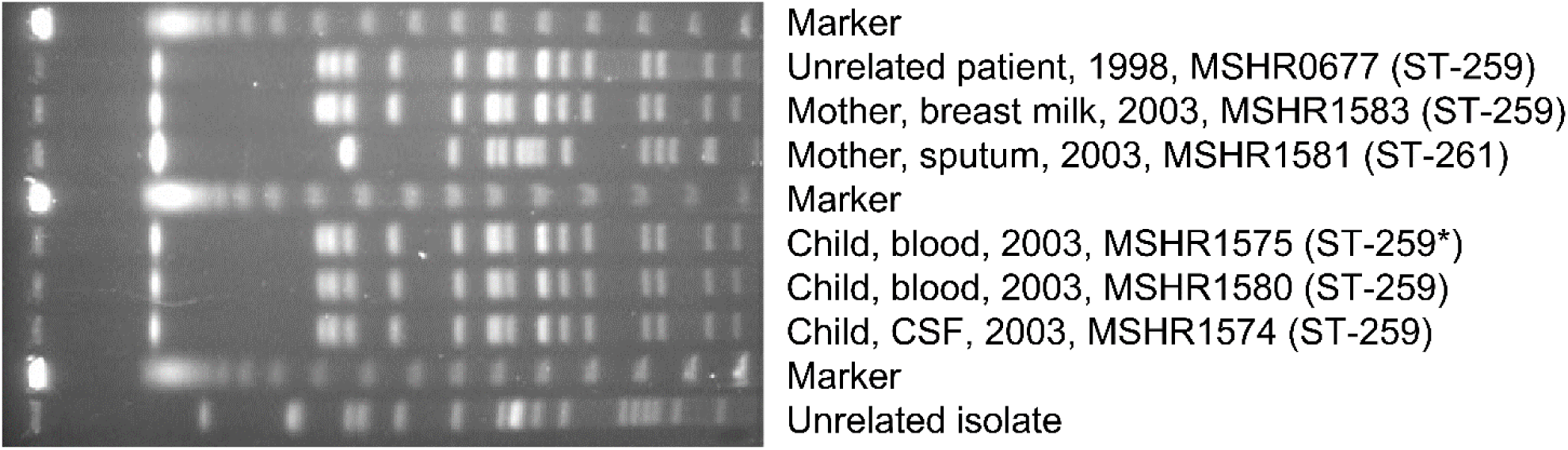
Pulsed-field gel electrophoresis analysis of mother and child isolates. *Isolate not subjected to whole-genome sequencing in this study due to subsequent culture destruction.

Prior studies have relied upon epidemiological and clinical observations [3, 11, 13, 14], often alongside gel electrophoresis-based genotyping methods [3, 11, 12], to examine cases of suspected *B. pseudomallei* transmission between human hosts. However, these genotyping methods lack the necessary resolution for definitive confirmation of such transmission events as they only assess a small fraction of the genome. As such, infections arising from independent environmental sources, or even from a single environmental point source as observed in outbreak scenarios [32, 36], cannot be ruled out using such lower-resolution methods. Consistent with the PFGE findings (Figure 1) [3], *in silico* MLST data showed strains from the mother’s sputum and breast milk matched the CSF- and blood-derived isolates retrieved from the child, with all isolates being ST-259 (Table 1). To obtain the most epidemiologically robust information from our WGS data, phylogenomic analysis of all mother-child ST-259 isolates was performed using a combined SNP-indel approach, which we have previously shown provides both higher resolution and a better fit with outbreak chronology compared with phylogenomic reconstruction using just SNPs [32]. This approach identified no SNP or indel differences between the mother and child ST-259 isolates (Figure 2A). Further comparative genomic analyses examining copy-number variants or larger deletions also failed to find any other genetic variation among the mother-child ST-259 isolates. Although there will always remain the possibility that the mother and child were infected from a single environmental point source, our collective clinical, epidemiological and genomic findings point strongly to ST-259 *B. pseudomallei* transmission from mother to child, with breastfeeding being the most likely route of infection. Our findings provide the strongest evidence presented to date that *B. pseudomallei* can transmit between human hosts. This finding raises clinical and biowarfare concerns, particularly in cases where a *B. pseudomallei* strain has developed acquired antimicrobial resistance (AMR) in one human host who subsequently transmits to another. Although acquired AMR in *B. pseudomallei* is relatively uncommon, there are myriad chromosomal mutations that can lead to clinically-relevant AMR in *B. pseudomallei* [37], leading to more challenging pathogen eradication [38]. While this phenomenon has not yet been documented, our study demonstrates that human-to-human transfer of an AMR *B. pseudomallei* strain is possible.

**Figure 2.**
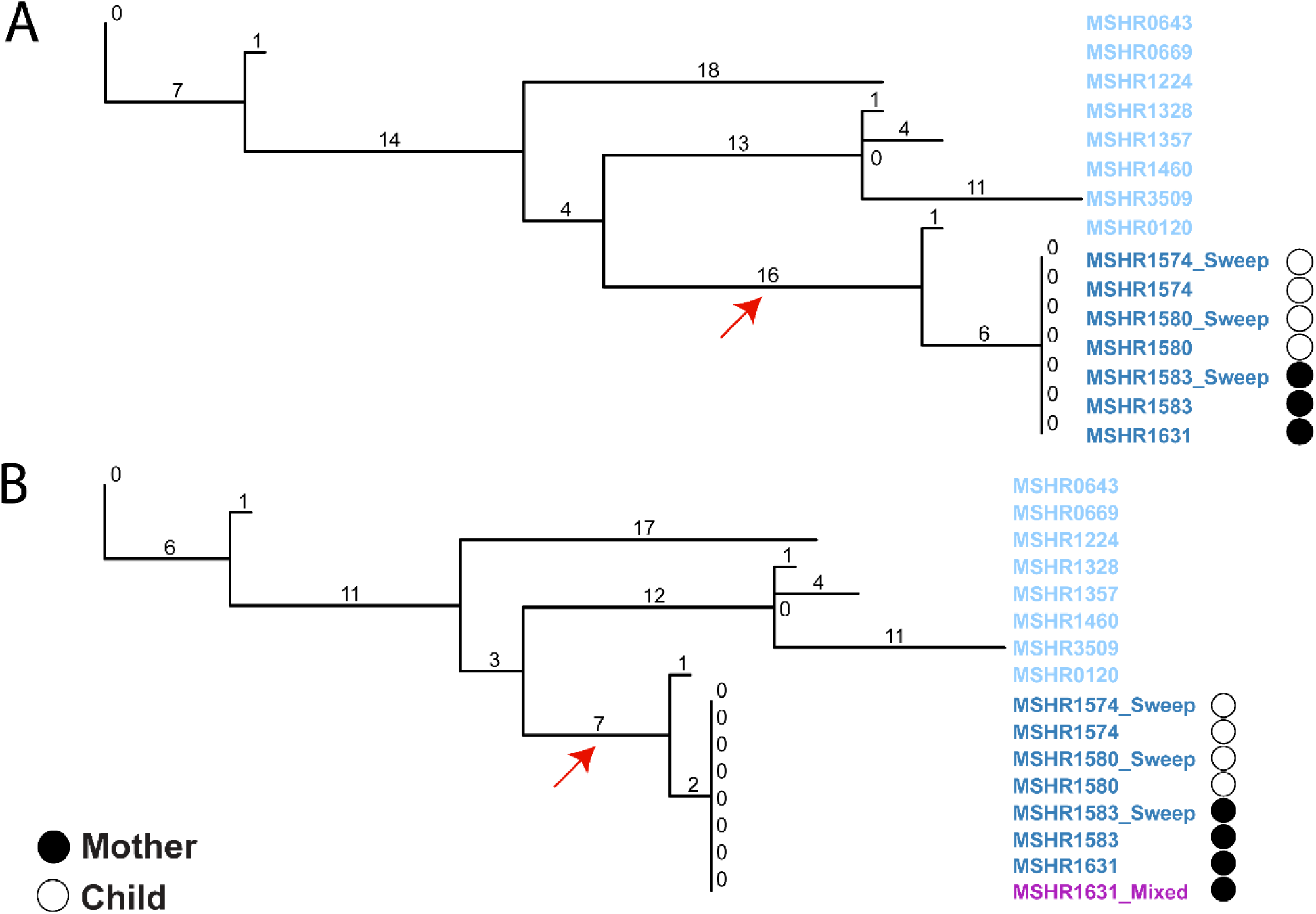
Maximum parsimony phylogenetic analyses of combined SNP-indel characters identified among *Burkholderia pseudomallei* ST-259 isolates, including mother and child isolates (dark blue). The MSHR1631_Mixed sample (purple) is a mixture of ST-259 and ST-261 at an 87%:13% ratio. (A) All ST-259 mother and child isolates were identical, with no observed SNP or indel differences. Mother-child isolates were most closely related to MSHR0120, a clinical ST-259 isolate from the same remote island that was collected in 1992. (B) The inclusion of a strain mixture (MSHR1631_Mixed; purple) from the mother results in the reduction of informative characters and branch collapse (red arrows).

To further understand ST-259 diversity on a broader scale, the ST-259 mother-child isolates were compared with seven temporally and geographically distinct clinical ST-259 isolates obtained between 1992 and 2009 from patients living in the Top End region of the Northern Territory. The mother-child clade was most closely related to MSHR0120, differing by seven variants (Figure 2). MSHR0120 was retrieved from a patient diagnosed with melioidosis 11 years prior who lived at the same remote locale as the mother and child. Additionally, minimal differences (between 36 and 45 variants) were observed between the mother-child clade and other ST-259 isolates, suggesting close relatedness of strains within this ST, but a clear difference between the mother-child cases and all other documented ST-259 cases in the Top End region. Taken together, these results provide further evidence for person-to-person *B. pseudomallei* transmission between mother and child.

Simultaneous infections with multiple *B. pseudomallei* strains have previously been reported [10, 39, 40]; however, the true rate of polyclonal *B. pseudomallei* infections is unknown. Polyclonality may increase the risk of neurological disease when one or more strains encode a *Burkholderia mallei bimA* (*bimA*_Bm_) genetic variant [41], and may cause issues with accurate point-source attribution in epidemiological investigations if polyclonality is not taken into account. Most clinical microbiological laboratories typically only select a single bacterial pathogen colony for further genotypic and phenotypic characterisation, which results in a considerable genetic bottleneck and the loss of strain mixtures from polyclonal clinical specimens. This shortcoming can be overcome using more time-intensive methods, such as the selection of multiple colonies for genetic analysis, sequencing of a ‘sweep’ of primary culture growth for further genetic characterisation, or by total metagenomic sequencing of the clinical specimen. Due to inherent ethical and technical issues with metagenomic sequencing of clinical specimens, we chose to genome-sequence culture sweeps and the individual colonies purified from them to identify putative *B. pseudomallei* strain mixtures in the mother and child clinical specimens. Consistent with the PFGE findings, *in silico* MLST and GATK HaplotypeCaller analysis of mother-child sweeps revealed that two distinct strains (ST-259 and ST-261) were found in one of the two sputa retrieved from the mother (MSHR1631_Mixed; Figure 3) but not in other primary sweep specimens from this patient (1 x sputum [MSHR1581_Sweep]; 1 x breast milk [MSHR1583_Sweep]), nor in the samples obtained from the child (1x CSF [MSHR1574_Sweep]; 1x blood [MSHR1580_Sweep]). WGS of single purified colonies from MSHR1631_Mixed and MSHR1581_Sweep confirmed that both ST-259 and ST-261 were present in this patient’s sputum specimens. Collectively, these results confirm that the mother had a simultaneous infection with two strains, adding to the documented polyclonal *B. pseudomallei* cases.

**Figure 3.**
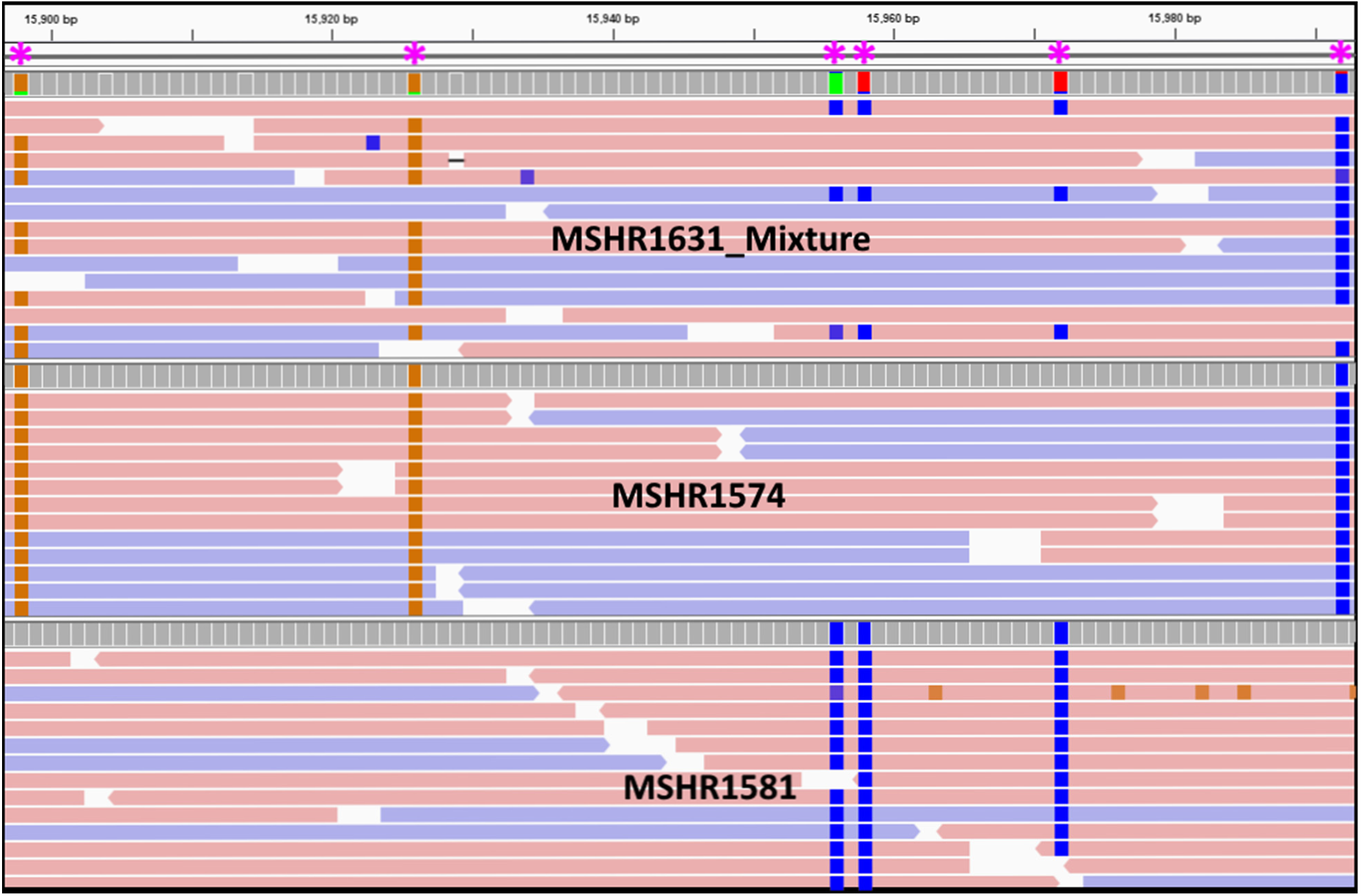
Example of ‘heterozygous’ (i.e. strain mixture) single-nucleotide polymorphism (SNP) calls at the sequence read level according to GATK HaplotypeCaller. Heterozygous SNP calls in MSHR1631_Mixed (ST-259 and ST-261) were parsed apart by comparing against homozygous SNP calls from MSHR1574 (ST-259) and MSHR1581 (ST-261). Horizontal bars represent forward (red) and reverse (blue) reads aligned against the MSHR1153 reference genome. Coloured boxes represent ‘heterozygous’ SNPs (asterisks).

To better understand this polyclonal infection from a bioinformatic standpoint, we first quantified the number of high-quality heterozygous SNPs in MSHR1631_Mixed. Haploid genomes such as bacterial genomes do not encode heterozygous SNPs; therefore, heterozygous SNPs are typically ignored by bacterial genome variant-calling software. The inclusion of heterozygous SNPs in an analysis of the mother-child isolates amongst a global dataset of *B. pseudomallei* genomes showed that MSHR1631_Mixed contained 12x the average number of heterozygous SNPs compared with all other mother-child samples (Figure 4). In total, 34,567 SNPs were identified in this sample, 47.8% of which were ‘heterozygous’. In contrast, an average of 29,914 SNPs were identified in the other nine mother-baby samples, of which only 5.15% were ‘heterozygous’. Next, homozygous SNPs identified in representative pure isolates (MSHR1574 for ST-259; MSHR1581 for ST-261) were used to identify the strain origin of each heterozygous allele from MSHR1631_Mixed SNPs. Using this simple method, 96% of heterozygous SNPs were matched to the correct strain. From these parsed data, we observed that 70% of heterozygous SNP read depths were within one standard deviation, with ST-259 dominant (87.1% of SNP read depths) and ST-261 present as a minor allelic component (12.9% of SNP read depths). No evidence of a tertiary strain was observed in the MSHR1631_Mixed when different ploidy settings were tested, indicating that no other strains were present.

**Figure 4.**
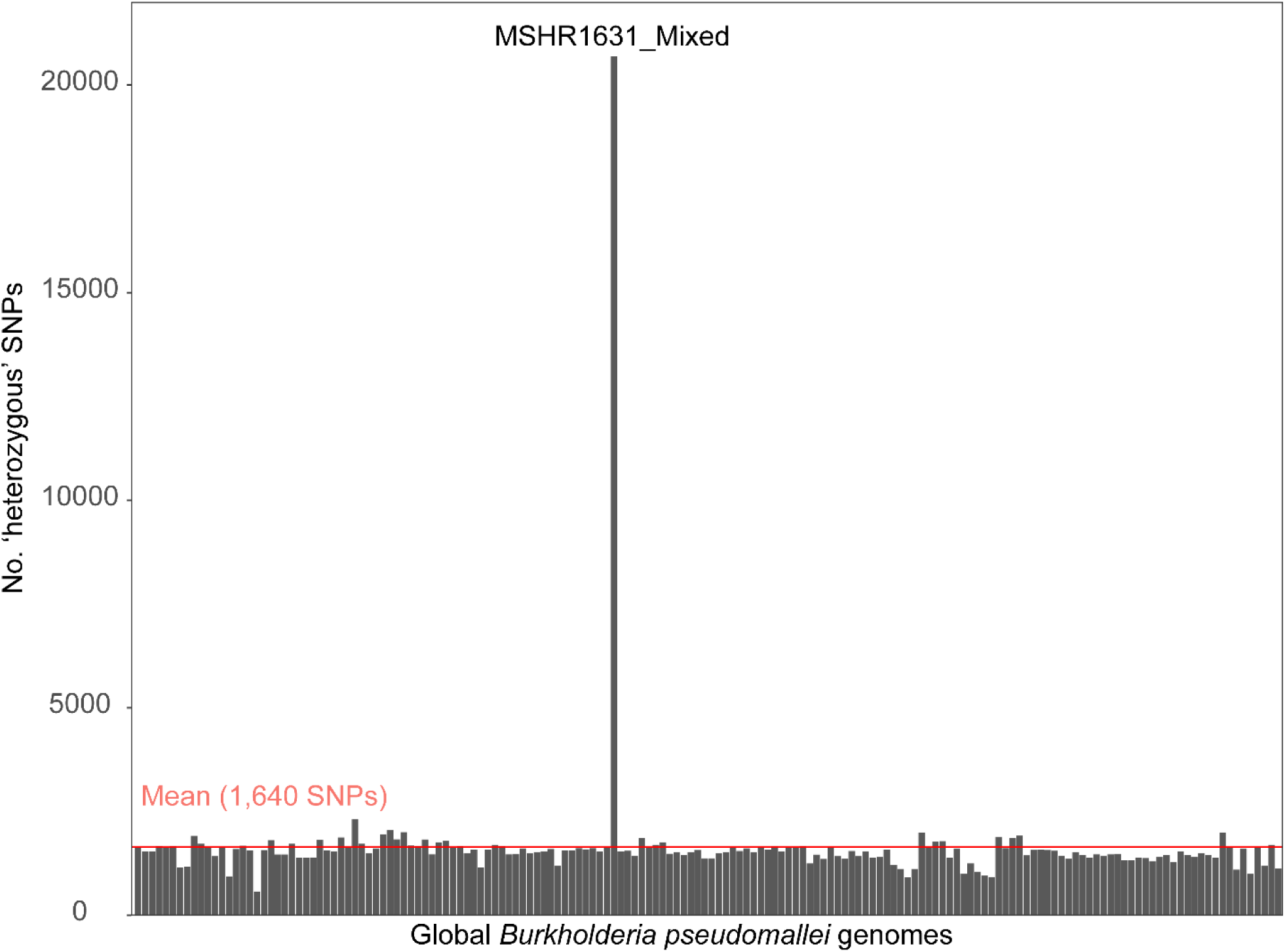
Quantification of ‘heterozygous’ (i.e. strain mixture) single-nucleotide polymorphism calls across all mother-child isolates and a global *Burkholderia pseudomallei* genome set. MSHR1631_Mixed contained 12x the mean number of ‘heterozygous’ calls according to the GATK UnifiedGenotyper, indicating the presence of a *B. pseudomallei* strain mixture in this sample. No other analysed genomes contained detectable mixtures.

The utility of SNP data derived from WGS to identify and study mixtures has been demonstrated in different diploid and polyploid organisms [42-44]. Current approaches in bacterial organisms include a database of known STs and proportion estimates of the bacterial population [45], which requires prior knowledge of the specific bacterial population or long-read sequencing [46], the latter of which is costly and error-prone when used in isolation. Bioinformatic solutions are available for ploidy inference of eukaryotic organisms [42-44, 47], which rely on the depth ratio of the two most abundant alleles sequenced for all heterozygous SNP positions across the genome (also referred to as ‘allele balance’). Such approaches assume SNP allele balances remain relative to each other; for example in a diploid sample, 50% of reads would support one allele while the other 50% support the other allele [42]. However, the allele balance assumption does not hold in bacterial mixtures, which may contain mixed ratios of any proportion. Despite this shortcoming, we demonstrated the feasibility of using SNP and read depth data to parse apart bacterial mixtures without any prior knowledge of the mixture composition. This approach relies on sequencing at a depth of ≥50x to ensure adequate sampling of a minor allelic component present at a ∼5-10% proportion. Such an approach is only suited for parsing apart mixtures of two strains. While the major strain is potentially identifiable in ≥3-strain mixtures, parsing apart minor components is a complex problem that remains unresolved using short-read data.

Finally, we investigated the effects of strain mixtures on phylogenomic reconstruction to determine whether the inclusion of even one mixture had unwanted effects on tree topology and phylogenetic inference. Phylogenomic analyses were performed with the ST-259 (Figure 2) and global (Figure 5) datasets, both with (Figures 2B and 5B) and without (Figures 2A and 5A) MSHR1631_Mixed inclusion. Tree comparisons identified two confounding issues in the trees containing MSHR1631_Mixed: branch collapse, and phylogenetic incongruence [48] that resulted in multiple instances of incorrect clade placement. In the ST-259 tree, the number of SNP-indel characters separating ST-259 isolates decreased from 35 to 21 variants (Figure 2B). In turn, the inferred relatedness between the mother-child ST-259 isolates and other ST-259 isolates was exaggerated due to the branch collapse (Figure 2B; red arrow). In the global dataset, branch collapse was also evident (Figure 5B). The cause of this branch shortening was the removal of all heterozygous SNPs from the dataset containing MSHR1631_Mixed, which reduced the total number of informative characters available for tree reconstruction by 18,051 SNPs when compared with the non-mixed phylogeny. Branch collapse was also evident in the global tree, whereby ST-261 isolates (MSHR1581_Sweep and MSHR1581; green text) incorrectly resided in the same clade as ST-259 (Figure 5B; asterisk). In contrast, the non-mixed dataset separated these two STs by approximately 20,000 SNPs, with clear separation of these clades (Figure 5A). Bootstrap values were of very high confidence across both trees at the ST-261 and ST-259 clades despite branch collapse and phylogenetic incongruence in the mixed dataset. Of further concern, the phylogeny containing MSHR163_Mixed caused incorrect geographic assignment of the Papua New Guinean clade, unexpectedly shifting its known grouping with Australian strains [9, 49] to the Asian clade; this incorrect placement received very high bootstrap support (Figure 5B). Reconstructing the global phylogeny sans MSHR1631_Mixed resolved both issues (Figure 5A).

**Figure 5.**
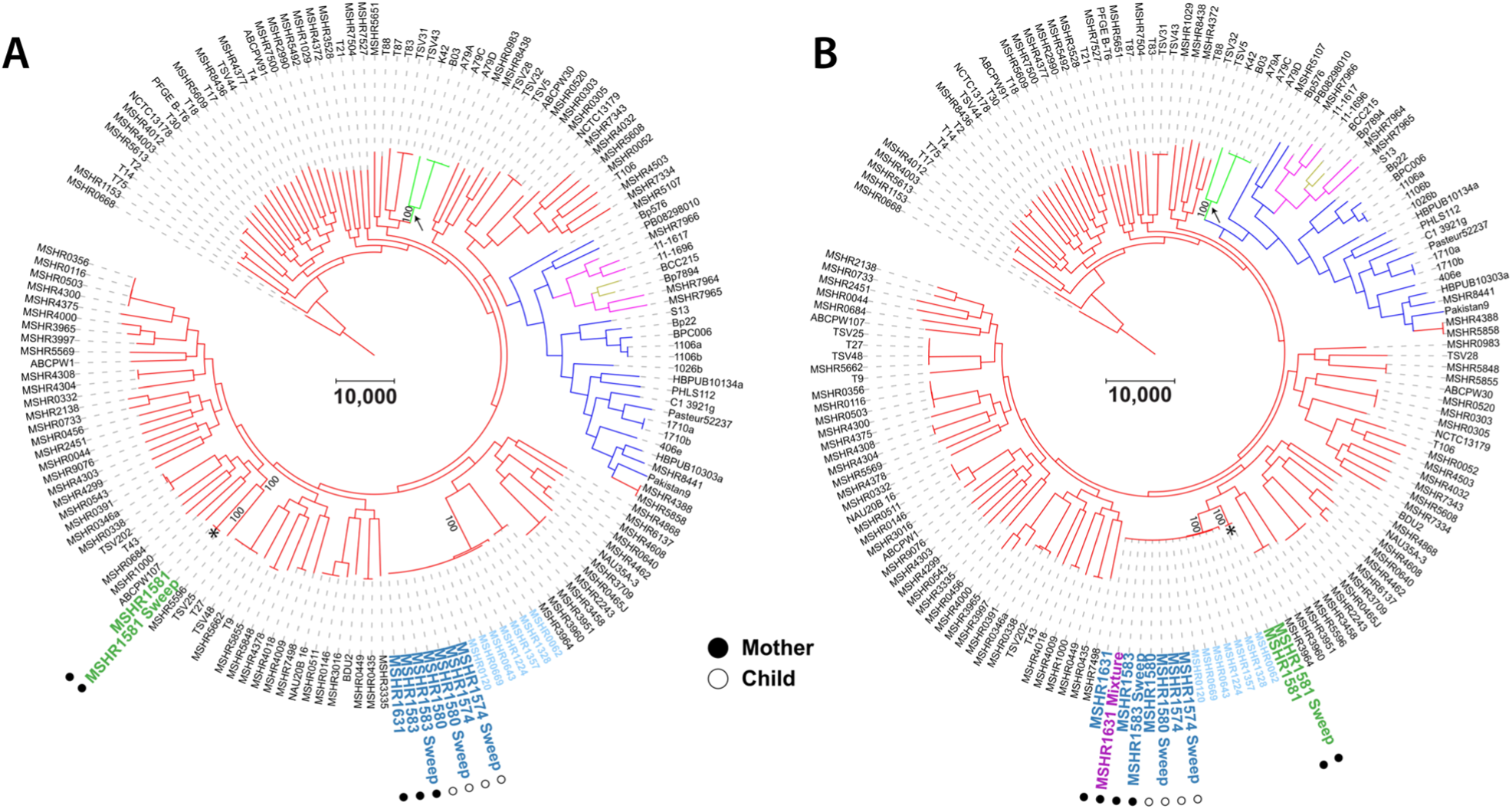
Global maximum parsimony phylogenetic analyses demonstrating the effects of strain mixtures on tree topology. Branch colours denote geographic origin of *Burkholderia pseudomallei* strains: red, Australian isolates, blue, Asian isolates; pink, African isolates; lime green, Papua New Guinean isolates; gold, South American isolates. (A) Exclusion of the mixed genome, MSHR1631_Mixed, results in correct topology and separation of STs 259 and 261 according to previous global *B. pseudomallei* phylogenies [9, 17, 31]; these two STs differ by >20,000 single-nucleotide polymorphisms (SNPs). (B) Inclusion of MSHR1631_Mixed greatly alters topology, leading to incorrect isolate and clade clustering, and collapsed branches in the clade containing MSHR1631_Mixed. Specifically, ST-261 isolates cluster incorrectly (asterisk) with ST-259, with branch collapse observed in this clade. The Papua New Guinean isolates are also incorrectly placed in this phylogeny (black arrows). The number of characters used to construct each tree differs by 14,503 SNPs (A: 207,209 SNPs; B: 192,706 SNPs). Dark blue text, ST-259 mother-child isolates; light blue text, geographically disparate ST-259 isolates; green text, ST-261 mother isolates.

The negative effects of strain mixtures on phylogenomic inference highlights the importance of strict quality controls throughout each stage of the experiment, especially during computational analysis. Bioinformatically, bacterial mixtures can be readily detected, as demonstrated in this study. However, standard practice in microbial variant calling pipelines is to report only homozygous variants for downstream analysis, with heterozygous SNPs typically ignored. Additionally, most phylogenetic reconstruction software treat heterozygous SNPs as missing or non-informative characters, even when encoded with IUPAC-ambiguous characters [50]. Our results provide unequivocal evidence that caution is needed in phylogenomic interpretation when dealing with potential strain mixtures. As these mixtures are not easily identifiable from phylogenetic analysis, it is prudent that microbial genomics studies include a mixture screening assessment of all genomes prior to variant calling and phylogenomic reconstruction to avoid removing phylogenetic informative characters, which can result in branch collapse or phylogenetic incongruence.

In conclusion, we demonstrate the utility of comparative genomics to both confirm human-to-human *B. pseudomallei* transmission and to identify simultaneous infection with multiple *B. pseudomallei* strains. Using a naturally-occurring mixed genome comprising two strains at an 87%:13% ratio, we describe an effective method to accurately identify and quantify such mixtures from WGS data, and highlight the confounding effects that even a single mixed genome can place on accurate phylogenomic interpretations for both closely related (e.g. single ST) and species-wide phylogenies. Our findings demonstrate the essentiality of assessing all microbial genome datasets for the presence of strain mixtures as a routine part of sequence data quality control. We strongly recommend that such mixtures be removed prior to phylogenomic analysis to avoid erroneous misinterpretations of strain relatedness.

## Author statements

BJC identified the transmission event, MM conducted specimen sample processing, PFGE, and DNA extractions. AA performed bioinformatic analysis with assistance and supervision from DSS and EPP. AA wrote the initial manuscript draft. DSS and EPP critically reviewed and edited the manuscript. BJC, MM, DSS, and EPP conceived of the study and obtained funding. All authors reviewed and approved the final manuscript.

## Data statement

All supporting data and protocols have been provided within the article.

## Data Bibliography

Accession numbers and references retrieved from Sarovich *et. al*. 2016 [31] for the 145 global *B. pseudomallei* isolate dataset is available on Figshare: https://doi.org/10.6084/m9.figshare.9840212

## Ethical approval

Ethics approval for this study was obtained from the Human Research Ethics Committee of the Northern Territory Department of Health and Families and the Menzies School of Health Research (HREC 02/38).

## Acknowledgements

We thank Vanessa Rigas (Menzies School of Health Research) for laboratory assistance.

